# Value coding by primate amygdala neurons complies with economic choice theory

**DOI:** 10.1101/2025.04.15.648888

**Authors:** Fabian Grabenhorst, Wolfram Schultz, Simone Ferrari-Toniolo

## Abstract

Primate amygdala neurons are implicated in coding the value of choice options for decision-making. However, it remains unclear whether amygdala value signals comply with the notion that decision makers maximize the value of rewards, as formalized by key axioms of economic theory. In particular, the continuity axiom of Expected Utility Theory (EUT) postulates that reward maximization is reflected by a trade-off between reward probability and magnitude. Our experiment tested this trade-off in two male macaque monkeys who subjectively ranked three gambles in choices, thus revealing their subjective value. Axiom compliance was defined by choice indifference between the intermediate gamble and a probabilistic combination of the other two gambles. We found that the animals’ choices between safe and gamble options reflected the integration of probability and magnitude into scalar values consistent with the continuity axiom. In a non-choice task, responses of individual amygdala neurons to separate probability and magnitude cues reflected the monkeys’ individual preferences: neuronal responses were lower for non-preferred gambles (relative to safe options), higher for preferred gambles and equal at subjective indifference between safe and gamble options. In a choice task, amygdala neurons integrated probability and magnitude into subjective values that reflected individual preferences. In some neurons, value signals transitioned to signals coding the monkey’s economic choices and chosen values. These findings identify amygdala neurons as substrates for encoding behavior-matching subjective values according to the continuity axiom of EUT, and for translating these values into economic choices.

**SIGNIFICANCE STATEMENT:** The amygdala—a brain structure in the temporal lobe—has long been associated with emotional responses. More recently, primate amygdala neurons have been implicated in economic decision-making. Here we show that amygdala neurons respond to visual cues indicating reward probability and magnitude in a manner that reflects the integration of these variables into subjective values. Importantly, these value signals complied with the notion that decision makers maximize the value of rewards, as formalized by key axioms of economic theory. Beyond valuation, some neurons processed subjective values into signals predictive of economic choices. Dysfunctional processing of values and choices by the amygdala could play a role in psychiatric and behavioral disorders affecting mental health, in which the amygdala is implicated.

## INTRODUCTION

Economic decision-making involves assigning subjective values to choice options and comparing these values between options. Although previous studies implicated neurons in specific brain areas in economic valuation and decision processes (Padoa-Schioppa and Assad, 2006; Schultz, 2006; Lau and Glimcher, 2008; Padoa-Schioppa, 2011; Grabenhorst et al., 2012; Chen and Stuphorn, 2015; Schultz, 2015; Lee and Seo, 2016; Grabenhorst and Schultz, 2021), fewer studies examined compliance of neuronal value signals with formally-defined aspects of economic choice theories (Padoa-Schioppa and Assad, 2008; Stauffer et al., 2014; Pastor-Bernier et al., 2019; Imaizumi et al., 2022; Yang et al., 2022; Ferrari-Toniolo and Schultz, 2023). A formal economic approach, combined with neurophysiological recordings, allows for the evaluation of abstract decision models at the individual-neuron level, supporting specific brain mechanisms that may underlie our choice behavior.

The primate amygdala, a nuclear complex in the medial temporal lobe, has been implicated in assigning value to stimuli and in economic decisions. Primate amygdala neurons encode the changing values of visual stimuli (Sanghera et al., 1979; Nishijo et al., 1988; Paton et al., 2006; Belova et al., 2007; Bermudez and Schultz, 2010; Costa et al., 2019), reward timing and contingency (Bermudez and Schultz, 2010; Bermudez et al., 2012), economic values and choices during individual decision-making (Grabenhorst et al., 2012; Jezzini and Padoa-Schioppa, 2020; Grabenhorst et al., 2023) and in social decision contexts (Chang et al., 2015; Grabenhorst et al., 2019b), probability and magnitude of expected rewards (Grabenhorst and Baez-Mendoza, 2025), self-determined plans to obtain future rewards through choice sequences (Hernadi et al., 2015; Grabenhorst et al., 2016), and formally defined decision computations translating values to choices (Grabenhorst et al., 2023). Consistently, amygdala signals in human neuroimaging studies reflect economic decision variables (Levy et al., 2010; Grabenhorst et al., 2013; Zangemeister et al., 2016; Kim et al., 2024). Previous studies showed neurons in other primate reward structures connected with the amygdala—the orbitofrontal cortex, dopaminergic midbrain and insula—code subjective values consistent with formally defined aspects of economic choice theories (Stauffer et al., 2014; Pastor-Bernier et al., 2019; Imaizumi et al., 2022; Yang et al., 2022; Ferrari-Toniolo and Schultz, 2023; Schultz, 2024; Ferrari-Toniolo et al., 2025). However, the value signals encoded by amygdala neurons have not yet been tested against formal requirements of economic choice theories.

Here we investigated whether the responses of amygdala neurons recorded in non-choice and choice situations are consistent with one of the axioms of Expected Utility Theory (EUT), the continuity axiom (von Neumann and Morgenstern, 1944). In choices between gambles (choice options varying in outcome probability and magnitude), the continuity axiom of EUT formalizes the integration of the reward components into a continuous subjective value quantity. According to the axiom, given three subjectively ranked gambles, the decision-maker should be indifferent between the intermediate gamble and a probabilistic combination of the other two. The existence of such an indifference point implies that no option is considered infinitely better than any other, which allows the definition of a finite, numerical subjective value for each gamble (Jehle and Reny, 2001). Importantly, the indifference point represents a quantification of the subjective value of the intermediate gamble in relation to the other two gambles. Continuity is a fundamental principle for economic valuation that is common to most modern economic choice theories, including developments of EUT addressing its empirical limitations (Samuelson, 1948; Weber and Camerer, 1987; Starmer, 2000).

By conducting tests according to the formalism of the continuity axiom of EUT, the present study defined a common, directly comparable subjective-value measure for both behavioral and neuronal data. We found that, by encoding behavior-matching subjective reward value, amygdala neurons are likely involved in reward maximization.

## METHODS

### Animals and ethical approval

Two adult male rhesus monkeys (Macaca mulatta) weighing 10.5 and 12.3 kg were used in the experiments. The animals had free access to standard laboratory-macaque diet before and after the experiments and, during recording periods, received their main liquid intake in the laboratory. All animal procedures conformed to US National Institutes of Health Guidelines. The work has been regulated, ethically reviewed and supervised by the following UK and University of Cambridge (UCam) institutions and individuals: UK Home Office, implementing the Animals (Scientific Procedures) Act 1986, Amendment Regulations 2012, and represented by the local UK Home Office Inspector; UK Animals in Science Committee; UCam Animal Welfare and Ethical Review Body (AWERB); UK National Centre for Replacement, Refinement and Reduction of Animal Experiments (NC3Rs); UCam Biomedical Service (UBS) Certificate Holder; UCam Welfare Officer; UCam Governance and Strategy Committee; UCam Named Veterinary Surgeon (NVS); UCam Named Animal Care and Welfare Officer (NACWO).

### Neurophysiological recordings

Experimental procedures for single-neuron recordings from the amygdala in awake, behaving macaque monkeys followed those described previously (Grabenhorst et al., 2012; Grabenhorst et al., 2019b). A titanium head holder and recording chamber (Gray Matter Research) were fixed to the skull under general anaesthesia and aseptic conditions. The amygdala was located based on bone marks on sagittal radiographs referenced to the stereotaxically implanted chamber (Aggleton and Passingham, 1981). We recorded the activity of single amygdala neurons from extracellular positions. We used standard electrophysiological techniques including on-line visualization and threshold discrimination of neuronal impulses on oscilloscopes. We aimed to recorded representative samples of neurons from the lateral, basolateral, basomedial and centromedial nuclei. We inserted a stainless-steel tube (0.56 mm outer diameter) to guide a single tungsten microelectrode (0.125 mm diameter; 1- to 5-MΩ impedance, FHC Inc.) through the dura. The microelectrode was advanced vertically in the stereotaxic plane with a hydraulic micromanipulator (MO-90; Narishige, Tokyo, Japan). Neuronal signals were amplified, bandpass filtered (300 Hz to 3 kHz), and monitored online with oscilloscopes. Behavioral data, digital signals from an impulse window discriminator, and analogue eye position data were sampled at 2 kHz on a laboratory computer with MATLAB (Mathworks Inc.) code. Analogue impulse waveforms were recorded at 22 kHz with a custom recording system and sorted offline using cluster-cutting and principal component analysis (Offline sorter; Plexon). We used one electrode per recording day and recorded between 1 and 10 neurons per day.

### Reconstruction of neuronal recording sites

After data collection was completed, the animals received an overdose of pentobarbital sodium (90 mg/kg iv) and were perfused with 4% paraformaldehyde in 0.1 M phosphate buffer through the left ventricle of the heart. We reconstructed the recording positions of neurons from 50-µm-thick, stereotaxically oriented coronal brain sections stained with cresyl violet based on electrolytic lesions (15–20 µA, 20–60 s, made in one animal) and lesions using cannulas placed to demarcate recording areas, by recording coordinates for single neurons noted during experiments, and in reference to brain structures with well-characterized electrophysiological signatures recorded during experiments (internal and external globus pallidus, substantia innominata) (DeLong, 1972). We assigned recorded neurons to amygdala nuclei based on reconstructed recording positions and a stereotaxic atlas (Paxinos et al., 2000) at three different anterior-posterior positions (figures in the paper show neuron locations collapsed over anterior-posterior levels).

### Non-choice (Pavlovian) task used for neurophysiological recordings

Two monkeys performed in a Pavlovian task with sequentially presented conditioned stimuli indicating the probability (as first cue) and magnitude (as second cue) of predicted liquid rewards under computer control (**Fig. 1A-C**). The monkey sat in a primate chair (Crist Instruments) in front of a horizontally mounted touch screen for stimulus display (EloTouch 1522L 15’; Tyco). Each trial started when the background color on the touch screen changed from black to gray. To initiate the trial, the monkey had to place his hand on an immobile, touch-sensitive key. Presentation of the gray background was followed by presentation of a central ocular fixation spot (1.3° visual angle). The animal was then required to fixate this spot within 4° for 500 ms and maintain fixation until reward delivery. The fixation spot was followed by presentation of a visual conditioned stimulus in the centre of the screen for 500 ms that indicated reward probability, drawn from a set of six stimuli (**Fig. 1C**). The probability stimuli were monochrome circular ‘sector’ stimuli; each sector stimulus consisted of two sectors distinguished by black-white shading at horizontal and oblique orientation with the amount of horizontal shading indicating the probability of obtaining the cued reward magnitude. Each stimulus predicted forthcoming reward with a specific probability of P = 0.37, P = 0.5, P = 0.63, or P = 0.75. Reward probability of P = 1 was cued by a gray square. This first cue period was followed by a 500-ms inter-stimulus interval which was followed by the second cue period, in which we presented a monochrome bar stimulus with the vertical position of the black horizontal bar against a white background indicating the magnitude of the predicted reward (drawn from a set of three magnitudes: 0.2, 0.4, 0.6 ml). A 500-ms inter-stimulus interval followed the second cue period before reward delivery. Reward delivery was followed by a trial-end period of 1,000 – 2,000 ms which ended with extinction of the gray background. The next trial started after an inter-trial interval of 2,000 – 4,000 ms (drawn from a uniform random distribution). A recording for a given neuron would typically last 90 trials.

**Figure 1.**
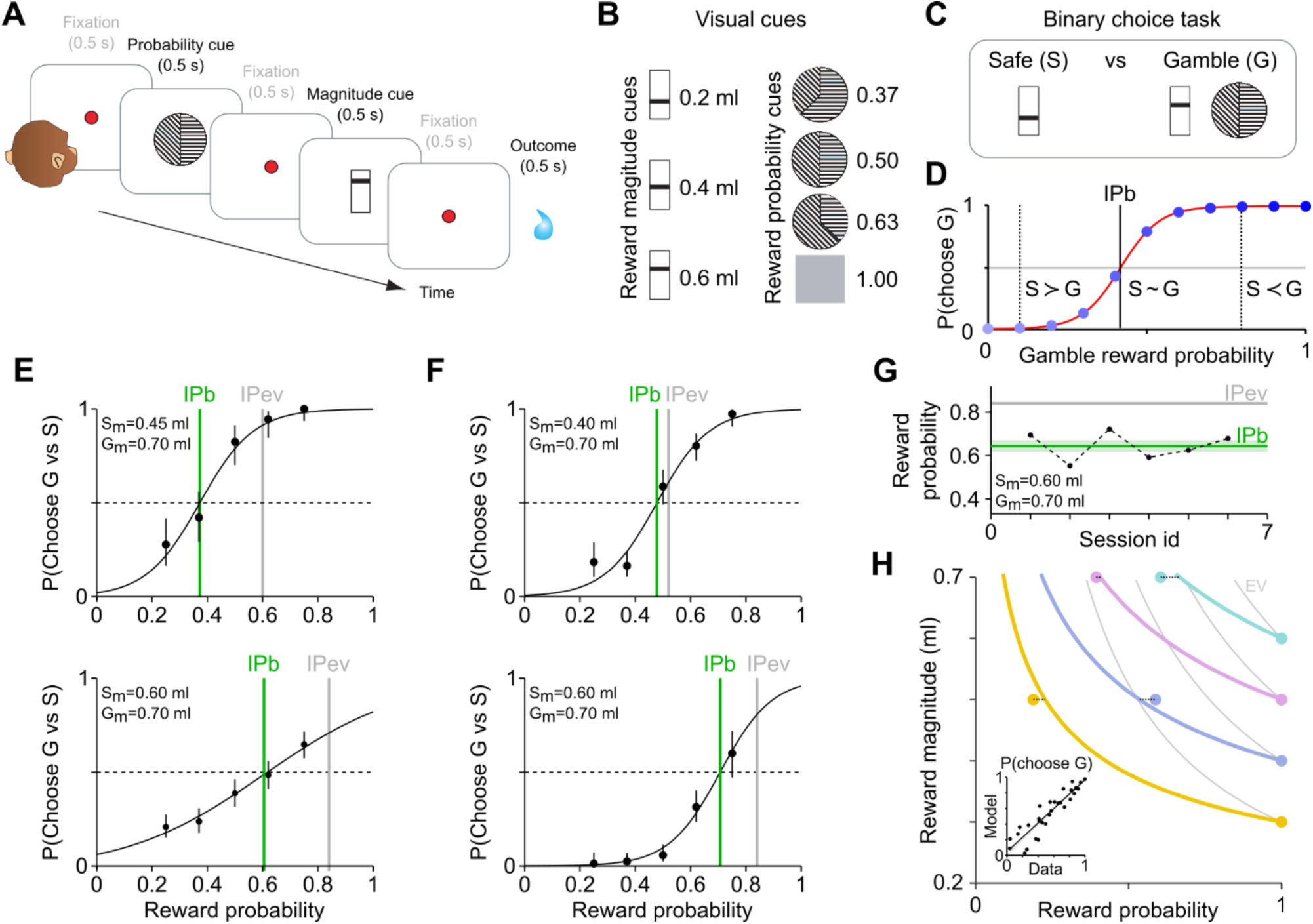
Experimental design and behavioral data. A) Trial structure. Pavlovian task for neural recordings: each trial started with the appearance of a red dot (fixation spot), which the monkey was required to fixate until reward delivery. A reward probability cue, a fixation spot and a reward magnitude cue appeared in successive 500 ms time intervals. After a further 500 ms fixation period, the reward outcome was delivered, contingent on the indicated reward magnitude and probability cues. B) Visual cues. The vertical position of a horizontal bar represented the reward magnitude (m) information, while a circular stimulus conveyed the probability (p) information. C, D) Indifference point estimation. Through binary choices between a fixed safe option and gambles with variable reward probability (C), we estimated the behavioral indifference point (IPb) as the reward probability for which the monkey was indifferent between the two options (D). We fitted a softmax function (red curve) to the proportion of gamble choices (blue dots) and identified the IPb as the gamble reward probability for which the softmax value was 0.5. This procedure was based on the continuity axiom of EUT, which states that a continuous variation of the reward probability should result in preferences shifting from the safe towards the gamble option, passing through an indifference point. The continuity axiom implies the existence of a continuously varying subjective value function. E-G) Elicited indifference points. Example behavioral indifference points (IPb, green line) elicited though binary choices between a safe option (magnitude Sm) and a gamble option (magnitude Gm) with varying reward probability (Gp). Dots: proportion of trials in which the gamble was chosen; vertical bars: binomial 95% confidence intervals. IPev (grey line): indifference point computed from objective reward values (i.e., probability Gp for which the gamble’s expected value (EV = Gm · Gp) equals the safe magnitude). An IPb lower than the IPev indicated a risk seeking attitude (i.e., gamble preferred to safe option with equal EV). Varying the safe magnitude from 0.45 ml (top) to 0.60 ml (bottom), resulted in a higher IPb, as expected from a monotonically increasing value function. Data from all sessions of monkey A (panel E) and monkey B (panel F); variability of IPb across different behavioral sessions (panel G), for one example test in monkey A. H) Indifference map. Choices between safe and gamble options with different Gm and Sm values resulted in a set of behavioral indifference points. Estimating an economic value function from behavioral choices (maximum likelihood procedure, see Methods), allowed us to compute a set of indifference curves (i.e., points with the same subjective value) which theoretically validated and interpolated the estimated IPb’s. Inset: significant correlation between measured and economic model-estimated choice probabilities (R^2^ = 0.84, P = 1.3e-12, N = 30). Data from monkey A.

Experimental conditions varied pseudo-randomly on a trial-by-trial basis. The specific reward probabilities and magnitudes were chosen based on pre-testing to ensure that the animals maintained high motivation during the task while at the same time providing sufficient variation to study neuronal activities related to probability, magnitude, expected value and risk. Together, the combination of the reward probability and reward magnitude cue on a given trial specified a probability distribution of possible reward magnitudes that could be delivered on that trial. Accurate reward prediction required the monkeys to combine information about the transiently cued reward probability and magnitude internally. A computer-controlled solenoid valve delivered liquid (juice) reward from a spout in front of the monkeys’ mouth. On each completed trial (without fixation breaks), the monkey received one of two outcomes: on ‘rewarded’ trials, we delivered a liquid reward corresponding to the cued reward magnitude in ml whereas on ‘non-rewarded’ trials, a small reward of 0.05 ml was delivered. We used small rewards rather than non-reward as we found that a small reward ensured that the animals maintained high motivation during recordings.

Possible errors in performance included failure to make contact with the touch-sensitive key before the trial, key release before trial completion, failure to fixate the central fixation spot at trial start or fixation break in the period between initial fixation and reward delivery. Errors led to a brief time out (3,000 ms) with a black background followed by trial repetition. We usually interrupted task performance after three consecutive errors. Fixation was continually monitored by the task program during all of these periods and fixation breaks resulted in an error trial. The animals were required to place their hand on a touch-sensitive key to initiate each trial and keep their hand in place on the key until trial completion.

Stimuli and behavior were controlled using custom MATLAB code (The Mathworks) and Psychophysics toolbox (version 3.0.8). The laboratory was interfaced with data acquisition boards (NI 6225; National Instruments) installed on a PC running Microsoft Windows 7.

### Behavioral choice task

We performed a separate choice task using the stimuli from the Pavlovian task to study the monkeys’ preferences for reward probabilities, reward magnitudes and associated expected value and risk levels and to confirm that the monkeys could use the information provided by these stimuli to make meaningful, reward-maximizing choices (preferring higher over lower reward probabilities and higher over lower reward magnitudes). The choice task was performed during the period of neurophysiological recordings but typically on separate testing days. In the choice task, reward magnitude was represented the same bar stimuli used in the main task and probability of reward was conveyed the same fractal or sector stimuli, presented adjacent to the bar stimulus. On each trial, the animal made a choice between two gambles, one of which was a ‘safe’ option, or ‘degenerate gamble’ (reward probability of P = 1, trial-by-trial varying reward magnitudes), presented randomly in left-right arrangement on the monitor. The safe option was cued only by a reward-magnitude cue, implying a reward probability of P = 1. For risky gambles, the cued reward magnitude could be obtained with the cued probability and a fixed small reward (0.05 ml) could be obtained with P = 1 – cued probability. Each trial started when the background color on the touch screen changed from black to gray. Trial start was similar to the main task. After 500 ms, the two choice options appeared on the touch monitor in left-right arrangement, followed after 750 ms by presentation of two blue rectangles below the choice options at the margin of the monitor, close to the position of the touch-sensitive key on which the animal rested its hand. The animal was then required to touch one of the targets within 1,500 ms to indicate its choice. Once the animal’s choice was registered, the unchosen option disappeared and after a delay of 500 ms, the chosen object also disappeared and a liquid reward was given to the acting animal. Reward delivery was followed by a trial-end period of 1,000 ms which ended with extinction of the gray background.

### Choice task used for neurophysiological recordings

The task has previously been described in detail (Grabenhorst et al., 2023). Two monkeys performed in a reward-based choice task with sequentially presented choice options under computer control (**Fig. 5A)**. On each trial, the animal made a choice between two sequentially presented options. Each option consisted of a visual ‘object’ (fractals, abstract images, photographs of natural objects such as flowers) presented in central position on the computer monitor overlaid by a small bar stimulus. Two different objects were associated with specific reward probabilities that varied across the testing session without notification. Different bar heights cued different reward magnitudes chosen randomly on each trial. To maximize reward, the animals were required to learn and track the (uncued) reward probabilities associated with the different objects and combine these probability estimates with the trial-specific cued reward magnitudes for the different objects. Reward probabilities varied in blocks of 15-40 trials and were pseudorandomly chosen for each object from the following set: 0, 0.15, 0.35, 0.5, 0.65, 0.75, 0.85, 1.0. Reward magnitudes varied randomly on each trial and were chosen from the following set: 0.25 ml, 0.4 ml, 0.65 ml. On each completed trial, the acting animal received one of two outcomes: on ‘rewarded’ trials, a liquid reward corresponding to the cued reward amount in ml was delivered whereas on ‘non-rewarded’ trials, a small reward of 0.05 ml was delivered. Each trial started when the background color on the touch screen changed from black to gray. To initiate the trial, the monkey was required to place his hand on an immobile, touch-sensitive key. Presentation of the gray background was followed by presentation of an ocular fixation spot (1.3° visual angle). On each trial, the animal was then required to fixate this spot within 4° for 500 ms. Following 500 ms of central fixation, a first choice cue (‘object’) and overlaid bar stimulus appeared centrally for 500 ms and were followed, after cue-offset, by a 500 ms inter-stimulus interval, which was then followed by a second choice cue and overlaid bar stimulus shown for 500 ms followed by another 500 ms inter-stimulus interval. The reward magnitude cue covered 18.75 percent of the underlying image. The two objects could have the same reward magnitude on a given trial, as determined by random permutation. We used new objects in each session. Following sequential presentation of these individual choice objects and overlaid bar stimuli, the two objects reappeared simultaneously on the left and right side of the monitor (determined pseudorandomly); importantly, the magnitude-bar stimuli did not reappear. Thus, the separate presentation of the first and second reward-magnitude cue, and their transient presentation during sequential viewing precluded simultaneous magnitude comparison. After 100 ms, the fixation spot disappeared, indicating that the monkey was no longer required to fixate the spot and was allowed to make his choice by fixating the object on the left or right for 500 ms. The monkey was allowed to freely look back and forth between the objects for 2,000 ms and in that period could make a choice at any time by fixating the chosen object for 500 ms. Once the monkey’s choice was registered, the unchosen object disappeared and after a delay of 500 ms, the chosen object also disappeared and a liquid reward was given depending on the scheduled reward probability and magnitude for the chosen option. Reward delivery was followed by a trial-end period of 1,000 – 2,000 ms which ended with extinction of the gray background. The next trial started after an inter-trial interval of 2,000 – 4,000 ms (drawn from a uniform random distribution). A recording session for a given neuron would typically last 150 trials.

Possible errors in performance included failure to make contact with the touch-sensitive key before the trial, key release before saccade choice, failure to fixate a choice object for 500 ms during the choice period, failure to fixate the central fixation spot at trial start or fixation break in the period between initial fixation and disappearance of fixation spot. Errors led to a brief time out (3,000 ms) with a black background and then trial repetition. Task performance was typically interrupted after three consecutive errors. The animals were required to fixate the fixation spot and the objects until the choice targets were presented in left-right arrangement. Fixation was continually monitored by the task program during all of these periods and fixation breaks resulted in an error trial. The animals were required to place their hand on a touch-sensitive key to initiate each trial and keep their hand in place on the key until trial completion.

Stimuli and behavior were controlled using custom MATLAB code (The Mathworks) and Psychophysics toolbox (version 3.0.8). The laboratory was interfaced with data acquisition boards (NI 6225; National Instruments) installed on a PC running Microsoft Windows 7.

### Task training

Following habituation to the laboratory environment and experimental set-up, we trained the animals over successive steps to drink liquid reward from the spout, place their hands on a touch-sensitive key and hold the touch key for increasingly longer periods to receive reward, to view different visual conditioned stimuli that resulted in reward delivery, to touch and choose between visual stimuli on a touch screen, to choose between visual stimuli based on fixed stimulus-associated reward probability or cued reward magnitude, to choose between visual stimuli under conditions of varying reward probability or magnitude, to choose between stimuli that varied in both reward probability and reward magnitude, to perform the task under head-fixation, to perform the task under gradually increasing visual fixation requirements including saccade choices. We progressed from task training to recording once the animals were implanted with recording chambers and when their performance had reached an asymptotic level. These training periods, including development of the tasks, lasted approximately 24 and 18 months for animals A and B.

### Eye data processing

We continuously monitored and recorded the animals’ eye positions using an infrared eye tracking system at 125 Hz (ETL200; ISCAN) that was placed next to the touchscreen. We calibrated the eye tracker before each testing during a fixation task. During recordings, accuracy of calibration of the eye tracker was checked and recalibrated if necessary.

### Using the continuity axiom for subjective value estimation

Expected Utility Theory (EUT) proposes four axioms that determine the maximization of utility (von Neumann and Morgenstern, 1944). Compliance with completeness (axiom I) and transitivity (axiom II) is necessary for consistently ranking all choice options. Compliance with continuity (axiom III) demonstrates that choices reflect a meaningful representation of numerical utility: the continuity axiom implies the existence of a value function. The independence axiom (axiom IV) defines the computation of expected utility from reward magnitudes and their probabilities. The four axioms of EUT define necessary and sufficient conditions for choices to be described by the maximization of subjective economic value: if the axioms are fulfilled, a subjective economic value can be computed from the reward components (magnitude and probability) for each choice option, and the decision maker behaves as if choosing the highest valued option. Violations of the independence axiom, reported in both human and animal subjects (Allais, 1953; Kagel et al., 1990; Ferrari-Toniolo et al., 2022), led to the development of non-expected utility theories, such as prospect theory (PT) (Kahneman and Tversky, 1979), which required a weakened form of the independence axiom while retaining the continuity axiom requirement.

The continuity axiom can be formally stated as:

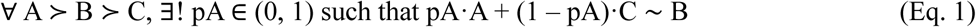

where A, B, C and AC are gambles, “≻” is the preference relation, and “∼” the indifference relation. In words, compliance with the continuity axiom requires to identify a reward probability pA at which choice indifference occurs between a fixed gamble B and a variable gamble AC. The variable gamble AC consists of a probabilistic combination of a higher valued gamble A (with probability pA) and a lower valued gamble C (with probability 1 - pA) (Eq. 1).

In both choice tasks used in the current study, the two offered choice options conformed with the continuity axiom definition. In the behavioral choice task (not used during neurophysiological recordings), we defined sure rewards (p = 1) as options A, B and C, resulting in binary choices between a fixed safe option (B) and a gamble with variable probability and fixed magnitudes (corresponding to the AC option in Eq. 1). In the choice task used for neurophysiological recordings, the B option was also a gamble, resulting in choices between a fixed gambles and a gamble with varying probability. The continuity axiom, with an extended testing scheme encompassing a broad set of reward magnitude and probability levels, constituted the foundation of the current study.

### Behavioral indifference points

The behavioral indifference point (IPb) represents a utility measure, being a numerical quantity associated with the subjective evaluation of reward B in relation to outcomes A and C. As a deterministic rule, the axiom assumes constant preferences over time. We interpreted the axiom in a stochastic sense, measuring preferences stochastically over repeated choices, which can potentially fluctuate over time. The IPb was computed according to a standard discrete choice model: we fitted a softmax function to the probability of choosing the AC option and identified the point for which the softmax’ value was 0.5, which corresponded to a 0.5 probability of choosing equally frequently each option, i.e., choice indifference (Figure 1C). The following softmax function was fitted to the choice data through non-linear least squares (Matlab function: nlinfit):

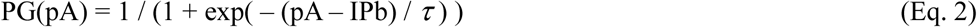

With PG representing the proportion of gamble choices, IPb corresponding to the probability pA resulting in equal preference for the two options, and *τ* as softmax “temperature” parameter, representing the steepness of the choice function (steeper for lower τ values), in analogy to the Boltzmann distribution in statistical mechanics. In alternative formulations, the softmax steepness is referred to as β (“inverse temperature” or “precision”), i.e., the reciprocal of the temperature parameter defined here.

### Behavioral indifference map

The indifference curves (ICs) represented in Figure 1H were elicited by fitting a parametric economic value function to the choice data. Following EUT, gamble values can be defined as the product of the corresponding utility and probability components. Using a parametric power function as utility function, we identified the best fitting parameters (softmax parameter, utility parameter) by maximizing the loglikelihood associated with the choice data, as described in detail in our previous behavioral study (Ferrari-Toniolo et al., 2021). As a goodness-of-fit measure, we correlated the probability of choosing the gamble option resulting from measured and modelled choices (Figure 1H inset). We then computed the values corresponding to finely spaced points (resolution 0.01 in both dimensions) within the tested range of reward magnitudes and probabilities. We defined the indifference curves as points with equal value (Matlab *contour* function), corresponding to four safe magnitude levels (0.3, 0.4, 0.5, 0.6 and 0.7 ml). The curves were plotted for visual comparison with the directly estimated IPs.

### Neuronal data analysis

We analyzed single-neuron activity by counting neuronal impulses for each neuron on correct trials in fixed time windows relative to different task events focusing on the following non-overlapping task epochs: 500 ms after onset of first cue (probability cue in the non-choice task, first choice option in the choice task), 500 ms after offset of first cue, 500 ms after onset of second cue (magnitude cue in the non-choice task, second choice option in the choice task), 500 ms after offset of second cue. We used fixed-window and sliding-window linear and multi-linear regression analyses to identify neuronal responses related to specific variables. For fixed-window analyses, we first identified task-related object-evoked responses by comparing activity in the cue and post-cue periods to a baseline control period (before appearance of fixation spot) using the Wilcoxon test (*P* < 0.05, Bonferroni-corrected for multiple comparisons). We then used multi-linear regression models to test whether neuronal activities were significantly related to specific task variables (P < 0.05, t-test on regression coefficient). We also used sliding-window multiple regression analyses with a 200-ms window that we moved in steps of 20 ms across each trial (without pre-selecting task-related responses). To determine statistical significance of sliding-regression coefficients, we used a permutation-based approach by performing the sliding-window regression 1,000 times using trial-shuffled data and determining a false-positive rate by counting the number of consecutive sliding-windows in which a regression was significant with *P* < 0.05. We found that less than five percent of neurons with trial-shuffled data showed more than ten consecutive significant analysis windows. Accordingly, we classified a sliding-window analysis as significant if a neuron showed a significant (*P* < 0.05) effect for more than ten consecutive 20-ms windows. Statistical significance of regression coefficients was determined using t-test; all tests performed were two-sided. We performed our regression analysis in the framework of the general linear model (GLM) implemented with the MATLAB function (*glmfit*). We used the following GLM for the non-choice task:

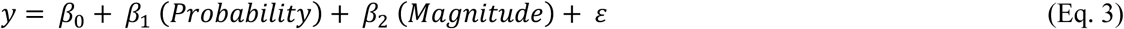

with y as the neuronal activity, *Probability* as reward probability and *Magnitude* as reward magnitude. This GLM served to identify probability-coding and magnitude-coding neurons in the non-choice task and to derive regression coefficients for Fig. 3C.

**Figure 2.**
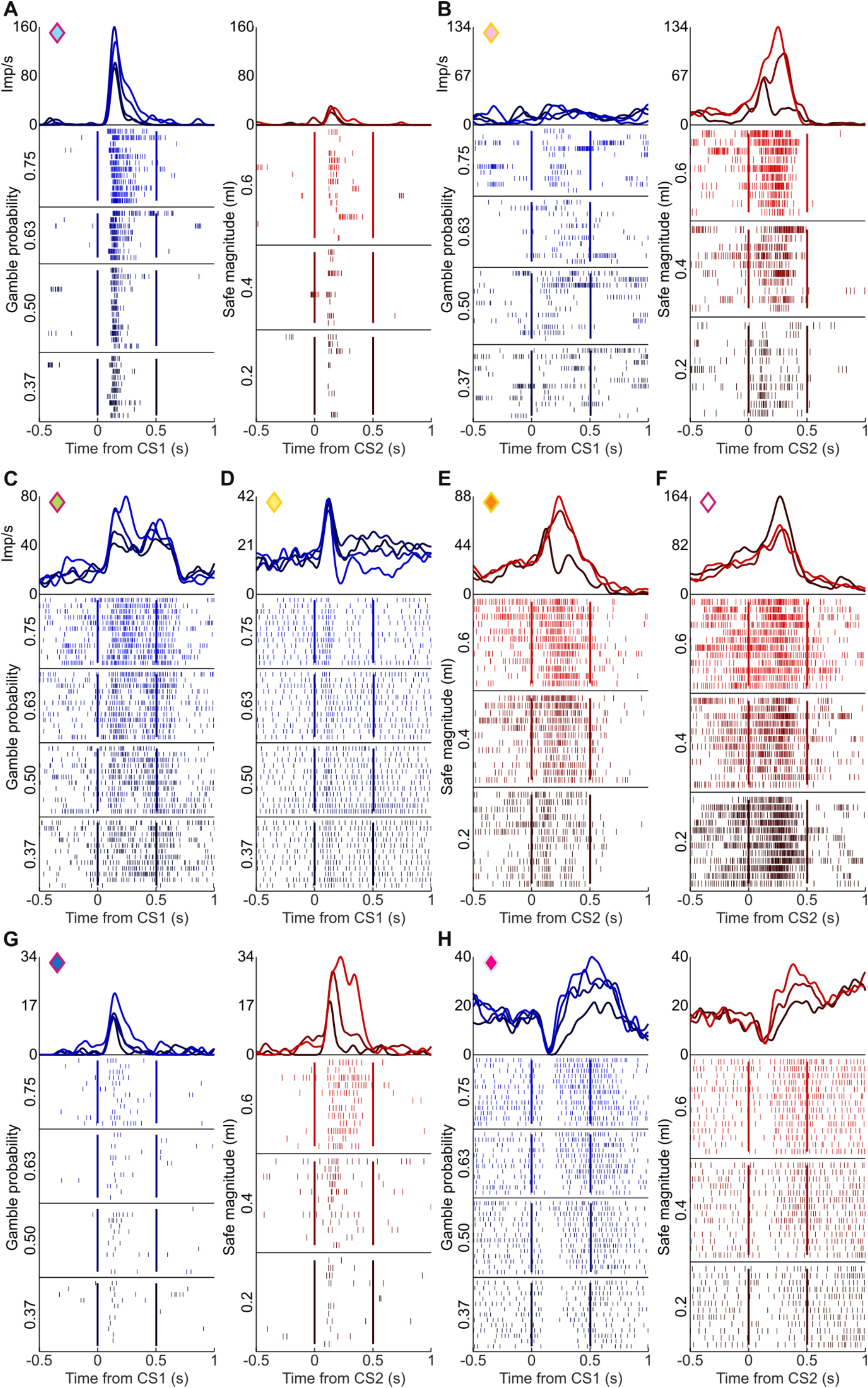
Coding of magnitude and probability in individual amygdala neurons. A, B) Example neurons coding either reward probability (p, panel A) or reward magnitude (m, panel B). Raster plot of action potentials (one trial per row) and spike density function (top; gaussian-convoluted activity rate, 25 ms kernel) for an individual neuron. Left: response to four options’ probabilities (Gp). Right: response to three safe options’ magnitudes (Sm). CS1: first conditioned stimulus, representing probability information; CS2: second conditioned stimulus, representing magnitude information (see Figure 1A, B). Diamond symbols identify each neuron’s anatomical location (see Figure 3A). C-F) Example neurons coding either reward probability (C, D) or reward magnitude (E, F). Neurons responded either proportionally (C, E) or inversely (D, F) to the coded variable (either reward magnitude or probability). G, H) Example neurons coding both reward probability and magnitude.

**Figure 3.**
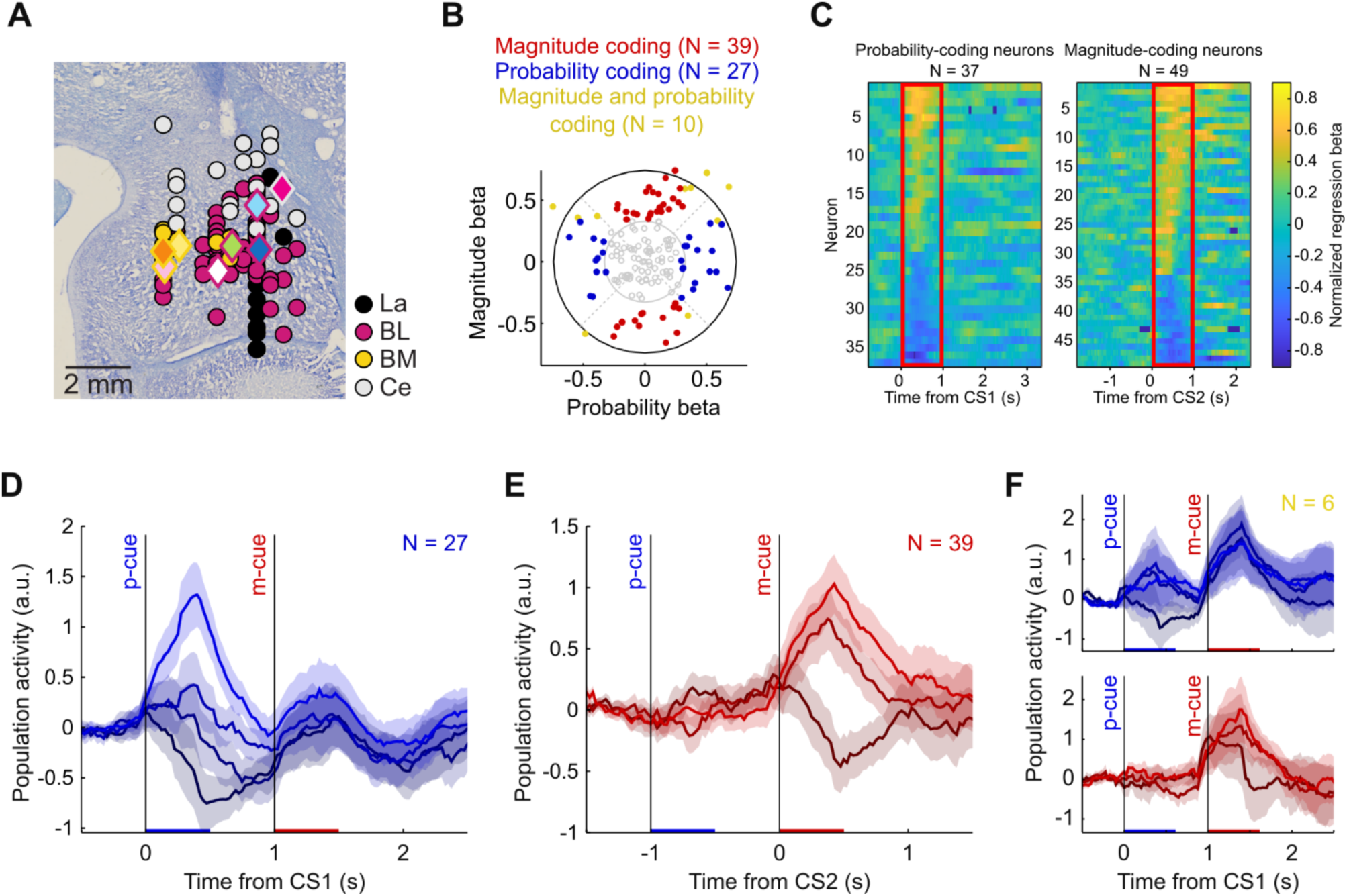
Coding of magnitude and probability in populations of amygdala neurons. A) Histological reconstruction of recorded neurons’ anatomical locations. Dots correspond to individual recorded neuron, fill colors identify the corresponding amygdala nucleus (La: lateral, BL: baso-lateral, BM: baso-medial, Ce: central). Diamond colors represent the example neurons reported in Figure 2 (fill) and identify the amygdala nucleus (outline). B) Time course of magnitude and probability coding. Neurons with significant probability-beta only (left) and magnitude-beta only (right) coefficients, sorted by beta values averaged within the 1 s post-cue time window (red rectangle). Both populations included neurons with significant positive and negative betas, corresponding to positive and inverse coding of the corresponding reward variable, respectively. C) Coding of magnitude and probability across the neuronal population. Normalized regression coefficients (betas) for magnitude and probability. Each dot represents a neuron with significant magnitude and probability betas (purple), significant magnitude beta only (red) or significant probability beta only (blue). Grey: no significant beta. D-F) Population responses. Average normalized population activity (area: SE) across the populations of magnitude-coding neurons (D) (3 magnitude levels: m = 0.2, 0.4, 0.6 ml), probability-coding neurons (E) (4 probability levels; p = 0.37, 0.50, 0.63, 0.75) and simultaneous magnitude-probability (same-sign) coding neurons (F). Neurons showing inverse coding of either magnitude or probability were rectified and included in the analysis.

We used the following GLM for analyzing neuronal activity during the first-cue period of the choice task:

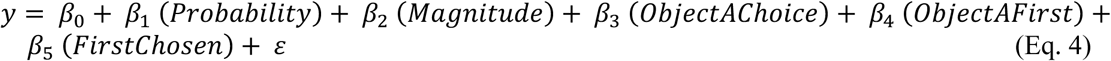

with y as the neuronal activity, *Probability* as reward probability, *Magnitude* as reward magnitude, *ObjectAChoice* as current-trial choice for object A (vs. object B), *ObjectAFirst* indicating whether object A was shown as first (vs. second) object on the current trial, and *FirstChosen* as choice of the first (vs. second) viewed object. This GLM served to identify probability-coding and magnitude-coding neurons in the choice task and to derive regression coefficients for Fig. 5C.

**Figure 4.**
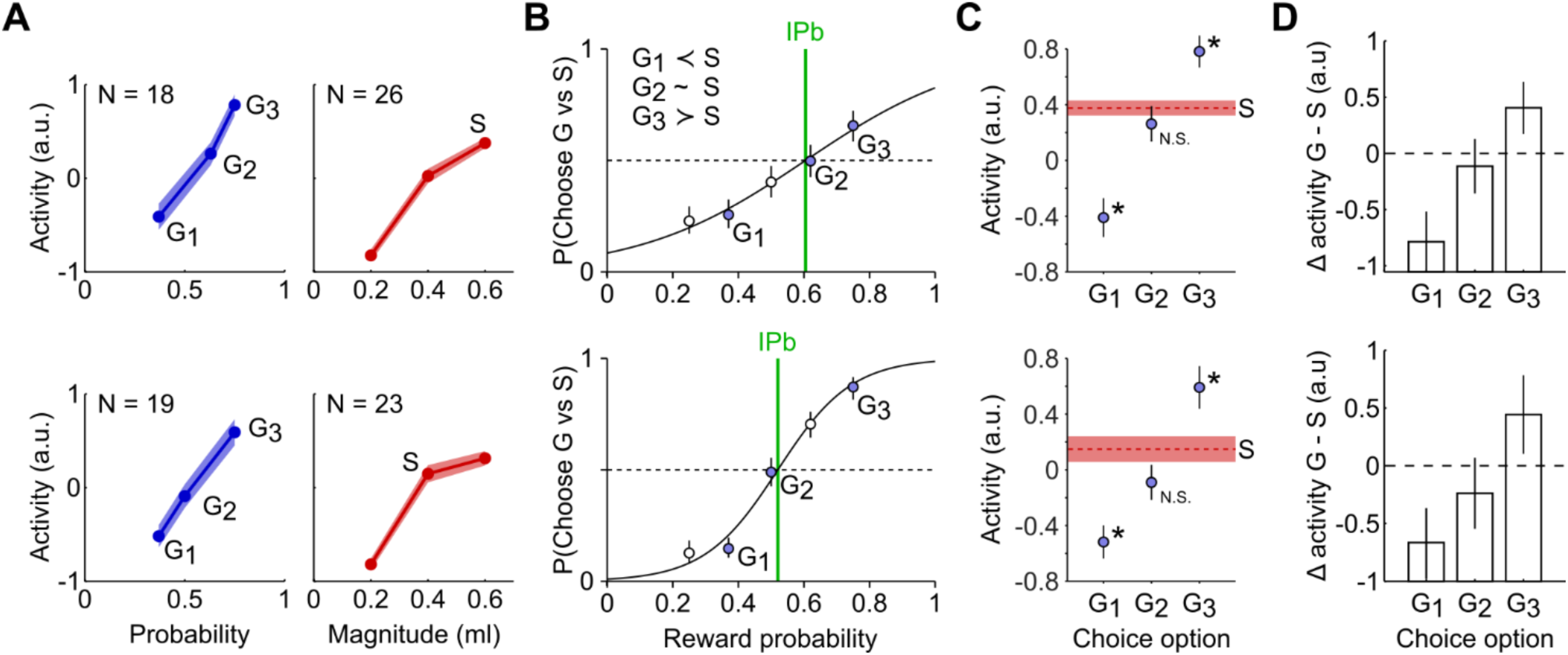
Match between behavioral and neuronal indifference points. A) Response to three gamble options (left) and three safe options (right) for the neuronal populations encoding reward probability and magnitude, respectively. Area: SE. S: safe option selected for subsequent data analysis. Data from monkey A (top) and monkey B (bottom). B) Behavioral preferences. Probability of choosing the gamble option against a fixed safe option, as a function of gamble probability. Monkeys had well defined preferences between each gamble (G_1_, G_2_ or G_3_) and the safe option S: G_1_ non preferred, G_2_ indifferent, G_3_ preferred. C) Population response to the selected options. Responses to the safe S option (dashed line: mean, area: SE) and to each of the three gambles (circles) were compatible with the behavioral preferences: response to S was significantly different from response to G_1_ and G_3_, while being not significantly different (N.S.) for G_2_. Statistical p-values (unpaired t test) for G_1_, G_2_ or G_3_, respectively: 1.0E-3, 0.35, 5.2e-7 (monkey A, top), 1.2e-2, 0.13, 5.9e-5 (monkey B, bottom). Asterisk, P < 0.05. D) Difference (Δ) in population activity between gamble and safe options. Compatible with a subjective-value coding signal, the activity difference was negative for G_1_, positive for G_3_ and non-significantly different from zero for G_2_. Bars: 95% confidence intervals.

**Figure 5.**
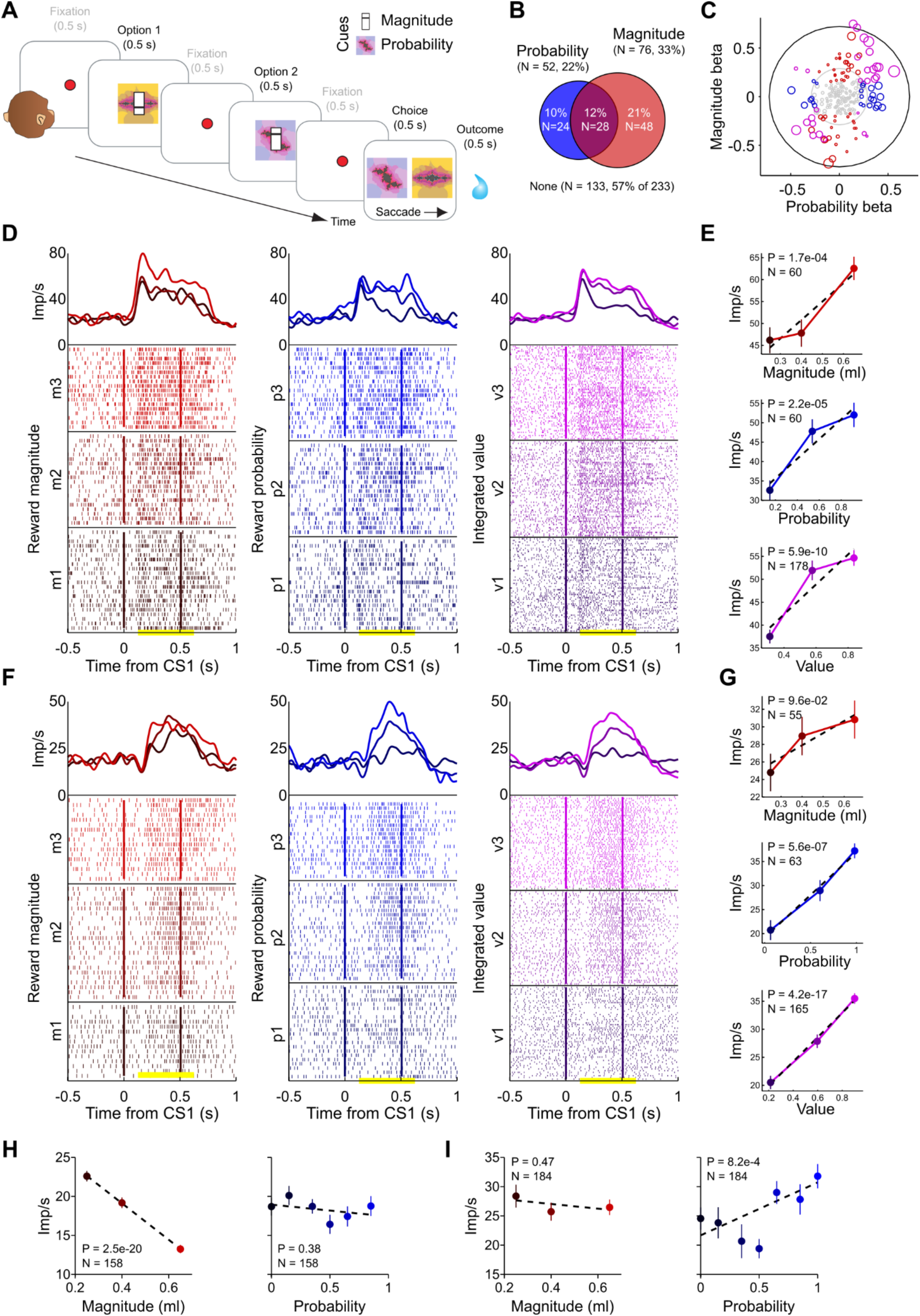
Integration of magnitude and probability information in individual neurons during choice. A) Choice task. Monkeys choose between sequentially viewed options based on reward probability and magnitude. Inset: Reward value derives from slowly changing, uncued reward probabilities and trial-specific, transiently cued magnitudes. B) Venn diagram representing neurons with significant regression coefficients for reward magnitude (red), probability (blue) or both (purple). Data from both monkeys. C) Regression beta coefficients for reward magnitude and probability. Colors identify significant betas for reward magnitude (red), probability (blue), both (purple) or none (gray). Circle size is proportional to the beta associated with the coding of integrated value. D) Response of an example neuron to the cue representing one choice option (raster plot and spike density function). The option’s reward magnitude (left) was explicitly cued (three levels), while the probability (center) was inferred from experienced rewards and its value was estimated through a reinforcement learning model. The model also computed a trial-by-trial estimate of the option’s integrated subjective value (right). The distribution of probabilities and subjective values were divided into terciles for data analysis, resulting in the three levels reported on the y axis (m_i_, p_i_, v_i_; i=1, 2, 3). The neuron’s responses significantly correlated with the cue’s reward magnitude, probability and integrated subjective value. E) Single neuron’s mean response rate ± SE (time analysis window: yellow line in panel A) to different magnitudes (top), probabilities (middle), and subjective values (bottom). Dashed line: linear regression. P: p value from correlation analysis. F, G) Response of a different example neuron encoding a cue’s reward magnitude, probability and integrated subjective value. Conventions as in panels D and E. H-I) Example neurons encoding exclusively reward magnitude (H) or reward probability (I).

We used the following GLM for analyzing neuronal activity using a sliding window of 200 ms moved in 20-ms steps across the first and second cue periods of the choice task:

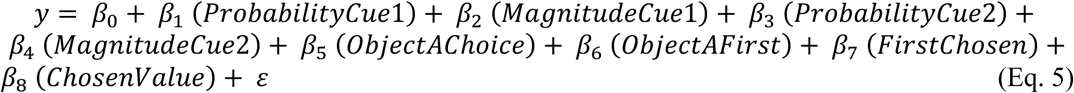

with y as the neuronal activity, *ProbabilityCue*1 and *ProbabilityCue*2 as reward probabilities of cues 1 and 2, *MagnitudeCue*1 and *MagnitudeCue*2 as reward magnitudes of cues 1 and 2, *ObjectAChoice* as current-trial choice for object A (vs. object B), *ObjectAFirst* indicating whether object A was shown as first (vs. second) object on the current trial, and *FirstChosen* as choice of the first (vs. second) viewed object and *ChosenValue* as the value of the chosen option on the current trial. This GLM served to identify probability-coding and magnitude-coding neurons in the choice task and to derive regression coefficients for Fig. 6D, E.

**Figure 6.**
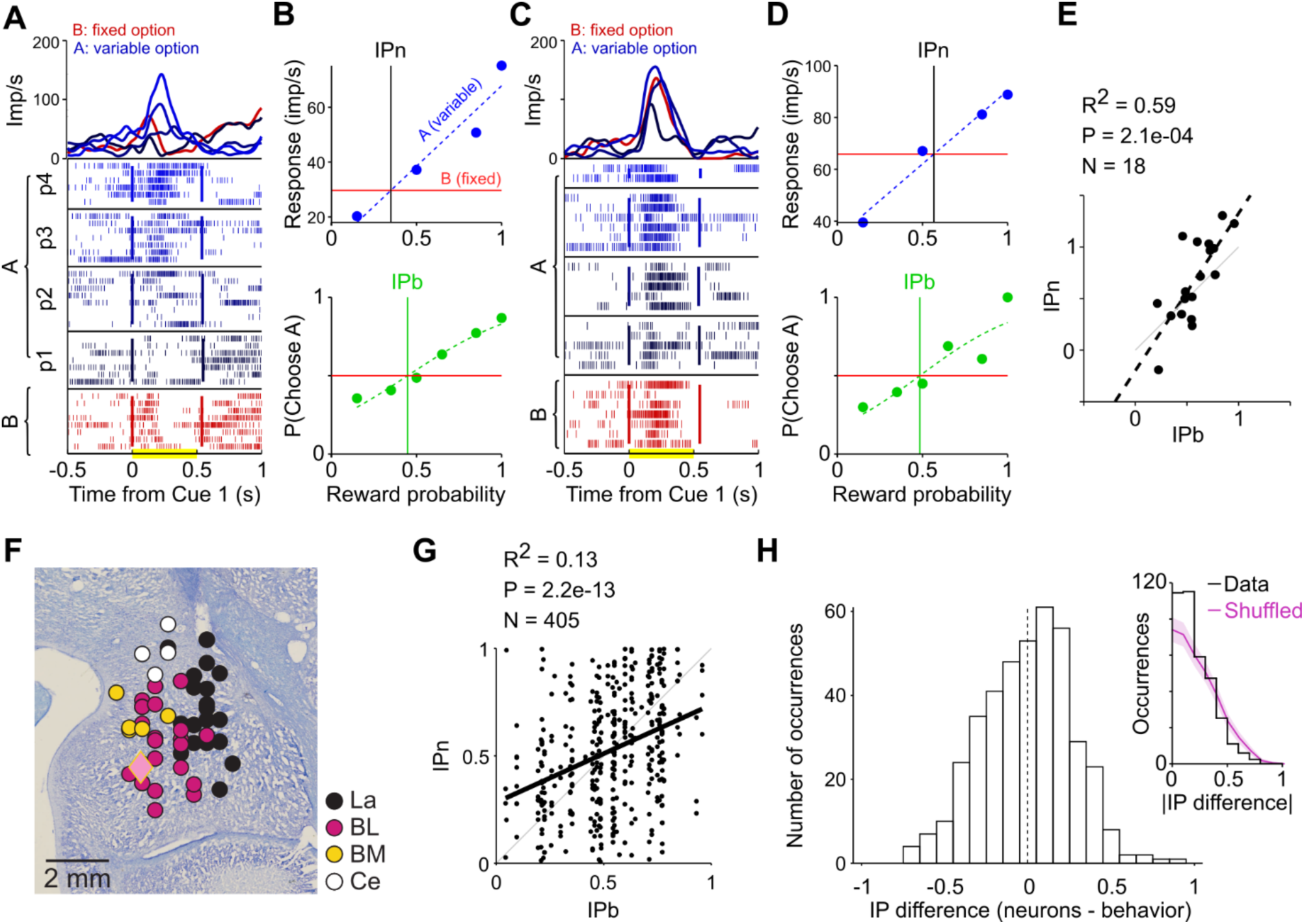
Behavior-matching neuronal subjective values compatible with the integration of magnitude and probability information. A) Neuronal responses of an individual neuron (raster plot and spike density function) to cues representing either a fixed choice option B (red; magnitude = 0.40 ml, probability = 0.15) or a variable choice option A (blue) with fixed magnitude (0.25 ml) and variable probability p_i_ (0.15, 0.50, 0.85 or 1.00). B) Neuronal (top) and behavioral (bottom) indifference points. Neuronal indifference point (IPn) was computed as the intersection between the linear fit (dashed line) of the mean responses to the variable option A (blue dots) and the mean response to the fixed B option (red line). Behavioral indifference point (IPb) was estimated via softmax fit to the probability of choosing the variable option A over the fixed option B (behavioral data from all sessions). C-D) Response of the same neuron as in panels A-B, for a different set of choice options (fixed B magnitude: 0.40 ml, probability: 0.85; variable option A magnitude: 0.65 ml). E) Correlation between the neuronal and behavioral indifference points (IPb and IPn respectively) across all tested choice options for example neuron (same neuron as in panels A-D). F) Histological reconstruction of the anatomical locations of the neurons modulated by both reward magnitude and probability in any task epoch (N = 61). We considered four 0.5 s task epochs: CS1, post-CS1, CS2 and post-CS2. Diamond symbol: example neuron shown in panels A to E. G) Correlation between neuronal and behavioral IP measures, across all neurons with significant IPn-IPb correlation and all tested combinations of choice options (N = 405). H) Distribution of differences between neuronal and behavioral indifference points (IPn – IPb) across the population of neurons with significant IPn-IPb correlation (N = 32), for all tested combinations of choice options (N = 405). Inset: distribution of absolute differences (|IPn - IPb|) computed from the collected data and from randomly shuffled data (shaded area: SD across 2,000 repetitions). The higher occurrence of lower differences in the original data compared to the randomly shuffled data (significant difference, Kolmogorov-Smirnov test, P = 8.5e-36, N = 405) supported the coding of behavioral-matching indifference points in amygdala neurons.

We adapted a method of classification of neuronal value responses based on the angle of regression coefficients (Wang et al., 2013; Tsutsui et al., 2016; Grabenhorst et al., 2023). In our case, this approach identifies probability-coding and magnitude-coding neurons by testing the statistical significance for a complete model that includes separate regressors for probability and magnitude. Using this method, a neuronal response was categorized as probability-coding or magnitude-coding if it showed a significant overall model fit (P < 0.05, F-test). For responses with significant model fit, we plotted the magnitude of the beta coefficients (standardized slopes) of the two probability regressors on an x-y plane. Following previous studies (Wang et al., 2013; Tsutsui et al., 2016; Grabenhorst et al., 2023), we divided the coefficient space into eight equally spaced segments of 45° to categorize neuronal responses based on the polar angle in this space of regression coefficients (Fig. 3B). We categorized responses as coding probability if their coefficients fell in the segments pointing toward 0° or 180° or as coding magnitude if their coefficients fell in the segments pointing toward 90° or 270°. We categorized responses as coding both probability and magnitude if their coefficients fell in the segments pointing toward 135° or 315° or in the segments pointing toward 45° or 225°.

### Normalization of population activity

To normalize activity from different amygdala neurons, we subtracted from the trial-specific impulse rate in a given task period the mean impulse rate and divided by its standard deviation (z-score normalization). We also distinguished neurons that showed positive relationships or negative relationships with a given variable, based on the sign of the regression coefficient, and sign-corrected responses with a negative relationship for plotting population activity. Normalized data were used for Fig. 3B-F, Fig. 4.

### Elicitation of neuronal indifference points

Mirroring the behavioral utility measure, we defined a neuronal subjective economic value measure for each tested set of A, B and C options. The neuronal indifference point (IPn) was defined as the gamble probability for which the neuronal response to the safe option matched the response to the gamble option. The neuronal indifference point (IPn) was defined as the gamble probability for which the neuronal response to the safe option matched the response to the gamble option. The IPn was computed as the intersection between two lines: the regression line of the neuronal activity for different gamble probabilities and the line representing the mean response to the safe option. To compute the IPn in individual neurons, responses to the two choice options were directly compared, thus no normalization of neuronal activity was required for this analysis.

## RESULTS

### Overview of the study

We investigated the encoding of reward probability, reward magnitude, and related subjective values in primate amygdala neurons across different tasks. We first examined the activity of amygdala neurons recorded in a Pavlovian, non-choice situation in which reward probability and magnitude were cued separately by sequentially presented visual stimuli (**Fig. 1**). Although the task design was based on a previous study (Grabenhorst and Baez-Mendoza, 2025), the neuronal data presented here are novel and recorded in conditions designed to test compliance with the continuity axiom of EUT. This study served to investigate whether separate probability and magnitude signals could provide a basis for the encoding of values consistent with the continuity axiom that defines conditions for maximizing reward. We then reexamined data from a previous investigation (Grabenhorst et al., 2023), in which amygdala neurons were recorded in a choice task with sequentially presented options varying in reward probability and magnitude. This study served to investigate whether individual amygdala neurons would combine probability and magnitude information into subjective values and related economic choices.

### Design and behavior

We recorded amygdala neurons in a Pavlovian task in which visual conditioned stimuli for reward probability and magnitude were shown sequentially on a given trial (**Fig. 1A**). This design allowed us to test if amygdala neurons would signal probability and magnitude consistent with principles of the continuity axiom of EUT. Each trial was defined by the combination of probability and magnitude cues drawn pseudorandomly from a set of seven distinct combinations (see Methods). These combinations were designed to test neuronal responses to probabilities and magnitudes close to the point of choice-indifference, assessed in a separate choice task (described below). Probability was cued by monochrome sector stimuli, whereas magnitude was cued by a bar stimulus (**Fig. 1B**). Using a Pavlovian, non-choice task allowed for a clear examination of neuronal probability and magnitude signals, as measured neuronal responses only depended on the cued stimuli but not on additional value-comparison and decision processes.

We used a separate choice task with the same stimuli to determine the monkeys’ subjective indifference points between combinations of probability and magnitude cues. The task was performed during the neuronal-recording period but mostly on separate testing days. The monkeys made binary choices between a ‘safe’ option defined by a fixed magnitude and a ‘gamble’ option defined by a combination of varying probability and fixed magnitude (**Fig. 1C**). By varying the gamble probability across trials and calculating the monkey’s probability of choosing the gamble at different magnitudes of the safe option, we determined the monkey’s indifference point (IP) from psychometric functions (**Fig. 1D**). This procedure was based on the continuity axiom of EUT, which states that a continuous variation of the reward probability should result in preferences shifting from the safe towards the gamble option, passing through an indifference point. The continuity axiom implies the existence of a continuously varying subjective value function, with the IP representing a quantitative measure of this subjective value. Comparing behaviorally determined indifference points (IPb) with hypothetical objective indifference points (IPev: reward probability for which the gamble’s expected value matched the safe magnitude), identified the monkeys’ risk attitudes. Monkeys were risk seeking if IPb < IPev (preferring the gamble even if its expected value was lower than the magnitude of the safe option) and risk averse if IPb > IPev (preferring the safe option even if its magnitude was lower than the expected value of the gamble).

**Figure 1E, F** shows choice data and measured indifference points in both monkeys elicited through the procedure described above. In line with the continuity axiom, when increasing the gamble’s reward probability, both animals gradually switched from preferring the safe option to preferring the gamble option. The resulting IPb’s varied when changing the safe and gamble magnitudes, suggesting that the animals considered both reward magnitude and probability information when making choices. The identification of an IPb indicated that magnitude and probability information were integrated into scalar values. The subjective nature of this value quantity was evidenced in the risk attitudes revealed by the IP measures. We found IPb’s lower than the respective IPev’s, implying a generally risk-seeking attitude in both animals. Our subjective-value measure was stable across testing sessions, consistently reflecting the animal’s risk attitude (Fig. 1G). We elicited IPb’s for different sets of safe and gamble magnitudes, which allowed us to estimate an economic value function from behavioral choices (maximum likelihood procedure, see Methods). The value function was then used to estimate indifference curves (i.e., points with the same subjective value) within a magnitude-probability space (Fig. 1H). This procedure validated our IPb measures, showing that the elicited indifference curves closely approximated the measured IPb’s across the full range of tested reward magnitude and probability levels (Fig. 1H, inset). As detailed in our previous extended behavioral study (Ferrari-Toniolo et al., 2021), this implies that subjective values can be described as the integration of mathematically defined reward magnitude and probability components.

Thus, the monkeys’ choices between safe and gamble options reflected the integration of reward probability and magnitude information into scalar subjective values, consistent with principles of the continuity axiom of EUT.

### Neuronal responses to reward magnitude and probability cues

We recorded the activity of 156 amygdala neurons in the Pavlovian task (75/81 neurons in animal A/B) across different amygdala nuclei: lateral (LA, 28 neurons), basolateral (BL, 85 neurons), basomedial (BM, 19 neurons), centromedial (Ce, 24 neurons). During the experiments, we sampled activity from about 300 neurons and typically recorded and saved the activity of those neurons that appeared to respond to any task event during online inspection of several trials. We aimed to identify task-responsive neurons but did not preselect based on particular response characteristics. This procedure resulted in a database of 156 neurons that we recorded and analyzed statistically.

Figure 2 illustrates the different observed response profiles of amygdala neurons, by showing the responses of ten amygdala neurons to cues indicating different levels of reward probability and reward magnitude. The neuron in Fig. 2A was recorded in the BL nucleus (cf. Fig. 3A) and showed a selective response to the probability cues that increased monotonically with the indicated reward probability; the neuron showed little response to reward-magnitude cues. By contrast, the neuron in Fig. 2B, recorded in the BL nucleus, showed the opposite activity pattern by responding with monotonically increasing activity to different cued reward-magnitude levels but showing little response to probability cues. Figure 2C-F illustrates further examples of probability-selective (Fig. 2C, D) and magnitude-selective neurons (Fig. 2E, F) with activities that either monotonically increased (Fig. 2C, E) or decreased (Fig. 2D, F) with increases in the encoded variable. Figure 2G**, H** shows two neurons with activity that increased with both reward probability and reward magnitude levels. Thus, individual amygdala neurons showed different types of activity patterns that coded the cued reward probability, reward magnitude or both variables. We next quantified the presences of these different neuron types in the population of recorded amygdala neurons.

Among 156 recorded amygdala neurons, 27 neurons (17%, 15/12 animal A/B) were classified as coding reward probability, 39 neurons (25%, 19/20 animal A/B) were classified as coding reward magnitude and 10 neurons (6%, 4/6 animal A/B) were classified as coding both probability and magnitude (Fig. 3B). Neurons were classified using a multiple-regression analysis based on the angle of probability- and magnitude-coefficients (Wang et al., 2013; Tsutsui et al., 2016; Grabenhorst et al., 2019a; Grabenhorst et al., 2023). Amygdala neurons coding reward probability and/or reward magnitude were found in all sampled amygdala nuclei (Fig. 3A, probability coding in LA/BL/BM/Ce: 4/15/4/4; magnitude coding in LA/BL/BM/Ce: 8/20/6/4; probability and magnitude coding in LA/BL/BM/Ce: 2/7/0/1). Responses of probability-coding neurons and magnitude-coding neurons were typically phasic, time-locked to the onset of the relevant cue, and showed positive and negative coding schemes in approximately equal proportions (Fig. 3C; probability-coding, positive/negative: 16/11; magnitude-coding, positive/negative: 26/13). The graded responses to different probability and/or magnitude levels were prominent in the aggregated population activity (Fig. 3D**-F**).

Thus, different groups of amygdala neurons coded the cued reward probability or reward magnitude and thereby signalled the two distinct components of subjective values in our task. We next examined whether these responses were consistent with key notions of the continuity axiom.

### Neuronal indifference points

Using the continuity axiom as a guiding principle, we tested whether amygdala neurons encoded reward information consistent with mathematically defined subjective values. To this aim, we analyzed the neuronal responses to different safe and gamble options for which we also measured the monkey’s behavioral preferences. Neuronal responses representing subjective values should reflect the monkey’s preferences, with higher response to a preferred option compared to the non-preferred option, and equal response for equally preferred options. In our continuity scheme, this implies that the gamble response, compared to the safe response, should be lower for non-preferred gambles, higher for preferred gambles and equal for equally chosen safe and gamble options.

Separately for the populations of probability-coding and magnitude-coding amygdala neurons, we pooled the neuronal responses to different gamble probabilities and safe-option magnitudes, respectively (Fig. 4A). To assess value coding in the neuronal population, we then selected three gamble options (G1, G2, G3) which were respectively behaviorally non-preferred, equally preferred, and preferred in relation to a fixed safe option (S) (Fig. 4B). This was done separately for each monkey, to account for their specific subjective evaluations. We found that the average responses to the selected safe and gamble options reflected the behavioral preferences. Compared with the responses to the S option, responses to G1 were significantly lower, responses to G2 were non-significantly different, and responses to G3 were significantly higher (Fig. 4C, D). This result was confirmed in both animals, reflecting their individual preferences.

These data show that the reward probability and magnitude components, separately coded by different groups of amygdala neurons, reflect individual preferences, suggesting the coding of subjective values for gamble and safe choice options.

### Coding of reward magnitude and probability during choice

So far, we examined the neuronal representation of reward magnitude and probability, separately cued in a non-choice, Pavlovian task. We next investigated the integrated coding of these fundamental reward variables in a choice task. This approach allowed us to identify neuronal signals reflecting behavior-matching subjective values resulting from the integration of reward magnitude and probability information and their translation to choices.

We recorded 233 amygdala neurons while monkeys performed a reward-based decision task (Grabenhorst et al., 2023). In each trial, two choice options were presented sequentially in two successive cue periods, followed by a side-by-side presentation of the same pair of options. The animals were required to integrate two value sources: tracking the slowly varying reward probabilities associated with each cue (different visual images), and combining them with trial-specific magnitudes cued by explicit bar stimuli (Fig. 5A, inset). Thus, each option’s reward magnitude was explicitly cued (three possible levels), while the probability had to be learned from experienced rewards. Magnitudes were cued transiently to encourage valuation and decision-making during sequential viewing. The monkey made a choice through a saccade towards one of the cues, receiving the corresponding reward outcome. All cue presentations (duration: 0.5 s) were separated by central fixation periods of 0.5 s (Fig. 5A).

We previously described that the monkeys’ choices in this task reflected the integration of reward probabilities and magnitudes for sequentially viewed options and that amygdala neurons encoded subjective values and related choices that reflected the integrated probability and magnitude information (Grabenhorst et al., 2023). Here, we analyzed the neuronal responses in relation to the separately varying magnitude and probability value components to test whether neuronal value signals complied with assumptions of the continuity axiom. In the first cue period, 24/233 amygdala neurons (10%) encoded reward probability but not magnitude, 48 neurons (21%) encoded reward magnitude but not probability, and 28 neurons (12%) were classified as encoding both probability and magnitude (Fig. 5B, C). The corresponding number of neurons encoding probability, magnitude or both probability and magnitude for the first post-cue delay period were 8 (3%), 26 (11%), 42 (18%); for the second cue period were 31 (13%), 29 (12%), 57 (24%), and for the second post-cue delay period were 7 (3%), 28 (12%), 38 (16%). The activity of the two amygdala neurons in Fig. 5D, E and Fig. 5F, G increased with both higher probability and higher magnitude levels at the first cue and accordingly were best explained in terms of integrated value coding. By contrast, the amygdala neuron in Fig. 5H encoded only the reward magnitude but not probability, while the amygdala neuron in Fig. 5I encoded only the reward probability but not magnitude.

Thus, in the value-based choice task, different amygdala neurons either integrated the reward probability and magnitude components into subjective value or selectively encoded on of these two value components. We next investigated whether these neurons’ activities were consistent with principles of the continuity axiom.

### Integration of magnitude and probability information during choice

To assess whether neuronal responses to the first cue were compatible with the coding of behavior-matching subjective values, we estimated and compared indifference points from behavior and neuronal data in the choice task. As with the Pavlovian task, this procedure was based on the continuity axiom of EUT, with one option (A) varying in probability (across blocks of trials), while the other option (B) was fixed. We analyzed the neuronal responses to options A or B in the first cue period (Fig. 6A, Eq. 4). A behavior-matching subjective-value coding neuron would respond equally to the A and B options that were equally chosen by the monkey. To test this notion, we calculated a neuronal indifference point (IPn) as the intersection between a linear fit of the mean responses to the variable option A and the mean response to the fixed B option. We compared the resulting IPn (Fig. 6B) with the corresponding behavioral IPb (Fig. 6C). We focused this analysis on neurons coding both probability and magnitude during the first cue period, to examine potential value-coding prior to value comparison and decision-making in the second cue period. Importantly, neurons used for this analysis were not preselected for coding subjective value but for coding both magnitude and probability.

In one example neuron (Fig. 6A**-D**), behavioral and neuronal indifference points appeared similar within the same set of A and B options (Fig. 6B, C). When varying the A and B options values, both IPb and IPn values appeared higher than those estimated for the first set of options (Fig. 6D). To statistically evaluate the neuronal-behavioral IP relation, we performed a correlation analysis across different A and B option sets (N = 18 option sets for this neuron). We found a significant correlation between IPb and IPn measures, suggesting that this example neuron encoded behavior-matching subjective values across the experienced reward magnitudes and probabilities (Fig. 6E).

In the population of amygdala neurons that responded significantly to both magnitude and probability (N = 28), 16 neurons showed a significant IPb-IPn correlation, while the average difference between IPb and IPn across neurons and IP measures (N = 170) was not significantly different from zero (t test, P = 0.72, N = 170, mean and SD: 0.0081 ± 0.29), suggesting coherent subjective-value coding in the population of amygdala neurons sensitive to both reward magnitude and probability. These results were confirmed when including three further task epochs of 0.5 s each (post-CS1, CS2 and post-CS2). We found N = 61 neurons responding significantly to both magnitude and probability in at least one of the four epochs. Among these neurons, which were distributed across different amygdala nuclei (Fig. 6F), we found 32 correlated neuronal and behavioral IP measures out of 74 total tests. A significant IPb-IPn correlation was confirmed in the full set of neurons and IP measures (N = 405) (Fig. 6G). The average difference between IPb and IPn was not significantly different from zero (t test, P = 0.81, N = 405, mean and SD: -0.0032 ± 0.27) and resulted in a better neuronal-behavioral match than randomly shuffled data (Fig. 6H).

These data show that a population of amygdala neurons represented the integration of reward magnitude and probability information during the choice task at the level of individual neurons by encoding a subjective-value measure that reflected the monkey’s preferences.

### Value-to-choice coding transitions in amygdala neurons

We next examined whether amygdala neurons that encoded value information were directly implicated in decision-making processes. We previously showed (Grabenhorst et al., 2012; Grabenhorst et al., 2019b; Grabenhorst et al., 2019a; Grabenhorst et al., 2023) that amygdala neurons implicated in economic decision-making exhibit dynamic coding patterns that transition from coding value (i.e., decision inputs) to coding choice (i.e., decision outputs). The amygdala neuron in Fig. 7A, B exhibited a value-to-choice transition by initially signaling the magnitudes of the first and second cues (Fig. 7A) and then signaling the monkey’s forthcoming choice for the first or second option on a given trial as well as the related chosen value (Fig. 7B). These dynamic coding transitions were evident in the regressions of the neuron’s activity on the different variables (Fig. 7C) and in phasic peaks of the coefficients of partial determination for each variable (Fig. 7D, Eq. 5). Dynamic value-to-choice transitions were expressed prominently in amygdala neurons that encoded the monkeys’ choices (N = 55, Eq. 5): in this subgroup of neurons, phasic signals related to the magnitude of the first and second option preceded signals related to the monkeys’ trial-specific choice and to the chosen value. Although individual neurons rarely showed the complex dynamic coding pattern shown in Fig. 7A, B (only 10/233 neurons (4%) coded all four variables), simpler coding transitions were more common: among 233 recorded amygdala neurons, 28 neurons (12%) coded the monkeys’ choice and the preceding magnitude of the first cue, and 21 neurons (9%) coded the monkeys’ choice and both the magnitude of the first and second cue.

**Figure 7.**
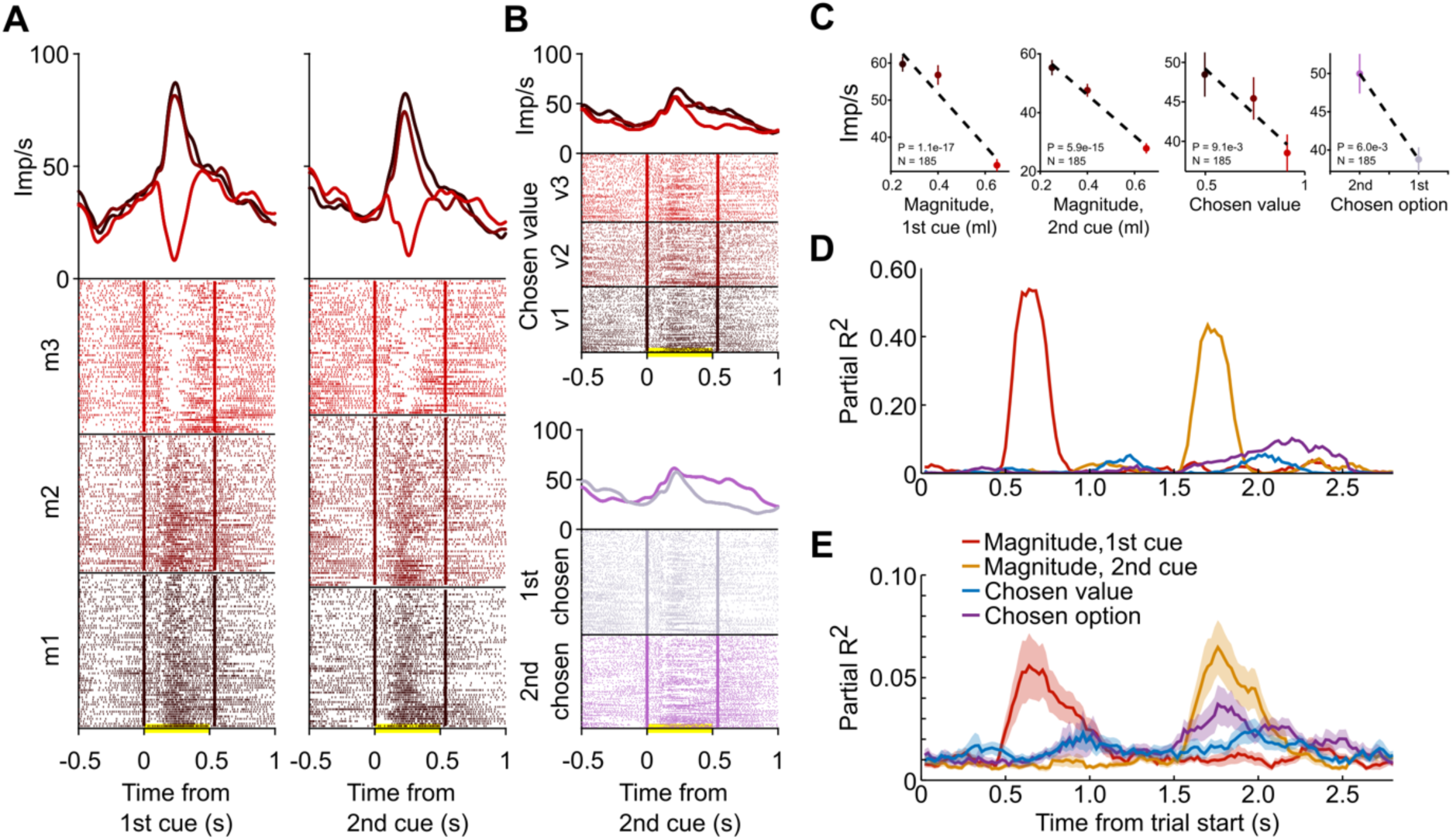
Neuronal value-to-choice transitions during decisions. A) Activity of one amygdala neuron coding the magnitude of the first (left) and second (right) choice option. B) Activity of the same neurons as in A) transitioned to coding the chosen value (top) and choice (bottom) on the current trial (right). C) Neuron’s mean response rate ± SE (time analysis window: yellow line in panels A) and B) to different magnitudes, chosen value, and choice. Dashed line: linear regression. P: p value from correlation analysis. D) Coefficients of partial determination (partial R^2^) from sliding-window multiple regression analysis of the neuron’s activity, showing periods of significant coding for magnitudes, chosen value and choice. E) Mean coefficients of partial determination (± SE) for 55 amygdala neurons identified as coding the monkeys’ choices in a sliding-window multiple regression analysis, showing population-coding of magnitudes of the first and second option, chosen value and choice.

Thus, the activity of amygdala neurons with phasic responses to reward-magnitude cues frequently transitioned to coding the monkeys’ choices and chosen values, consistent with involvement of these neurons in economic decision processes.

## DISCUSSION

These results show that primate amygdala neurons encode the basic components of economic value, reward magnitude and probability, in compliance with formal tenets of economic theory that deal with reward maximization. According to the continuity axiom of EUT, a decision-maker should be indifferent between the intermediate of three subjectively ranked gambles and a probabilistic combination of the other two. The compliance of the monkeys’ behavioral choices with the continuity axiom is consistent with the integration of reward magnitude and probability into a scalar value quantity. This integration represents the continuous tradeoff between reward components (a decrease in magnitude is compensated by an increase in probability, and vice versa), as shown in the behavioral indifference map (Fig. 1). The robustness of these conclusions relied on testing the axiomatic rule with a sufficient range of magnitude and probability levels. When tested in neurons, compliance with the continuity axiom reflected the integration of magnitude and probability information into a value signal. By itself, this result does not demonstrate encoding of subjective values, as it may simply reflect the monotonic neuronal responses to both reward magnitudes and probabilities (Fig. 2, 3, 5). However, by comparing the behavioral and neuronal subjective-value measures (i.e., the indifference points based on the continuity axiom), we ensured that the neuronal value code reflected the individual animal’s subjective evaluation of the choice options, in both no-choice (Fig. 4) and choice situations (Fig. 6). Single-cell and population signals translating these subjective evaluations into choice predictions and chosen-value signals (Fig. 7) demonstrate that amygdala neurons encode key variables underlying economic choice in a manner suitable for maximizing reward.

Although previous studies identified neurons encoding subjective values in the amygdala (Grabenhorst et al., 2012; Hernadi et al., 2015; Costa et al., 2019; Grabenhorst et al., 2019b; Jezzini and Padoa-Schioppa, 2020; Grabenhorst et al., 2023), the orbitofrontal cortex (Padoa-Schioppa and Assad, 2006, 2008; Pastor-Bernier et al., 2019; Imaizumi et al., 2022; Ferrari-Toniolo and Schultz, 2023), and other brain structures (Samejima et al., 2005; Lau and Glimcher, 2008; Padoa-Schioppa, 2011; Cai and Padoa-Schioppa, 2012; Lak et al., 2014; Stauffer et al., 2014; Schultz, 2015; Tsutsui et al., 2016; Costa et al., 2019; Grabenhorst et al., 2019a; Yang et al., 2022; Schultz, 2024), only two previous studies examined compliance of neuronal value signals with the continuity axiom of EUT (Ferrari-Toniolo and Schultz, 2023; Ferrari-Toniolo et al., 2025). The present results contribute to the emerging picture of a neuronal value-based choice mechanism compatible with axiomatic economic principles, distributed across core reward-related primate brain regions, including the amygdala, the orbitofrontal cortex and the midbrain dopamine area.

In our non-choice task with separately cued reward magnitude and probability components, the majority of amygdala neurons encoded only one of the two reward components. This might reflect subjective value coding for specific categories of choice options (in this case, safe and gamble options). Alternatively, it could reflect the coding of only one reward component, a feature that would be required for integrating magnitudes and probabilities into a single scalar quantity. As we only used one reward magnitude for the gamble option in the no-choice task, further tests are necessary to discriminate between these two hypotheses, by simultaneously varying both reward magnitude and probability of the gamble options. Another possibility is that amygdala neurons encode reward and decision-variables in a context-sensitive manner. For example, previous studies showed that neuronal coding of valuation and decision variables in a sequential reward-saving task was for many amygdala neurons specific to free-choice (compared to forced-choice) contexts (Grabenhorst et al., 2012; Hernadi et al., 2015; Grabenhorst et al., 2016). Our recent study on amygdala neurons in a non-choice task, using different task parameters, showed that few neurons integrated probability and magnitude into expected value without explicit requirement for decision-making (Grabenhorst and Baez-Mendoza, 2025). Our present results on amygdala neurons in non-choice and choice situations show that amygdala neurons do integrate both value components into subjective value signals in choice situations. Thus, neuronal integration of reward parameters into decision variables in the amygdala appears sensitive to context, i.e., the presence of decision-making requirements.

EUT defines the exact computation for combining reward components into scalar subjective values. According to EUT, the subjective value coincides with the utility associated to a choice outcome weighted by its probability of occurrence. The fourth axiom of EUT, the independence axiom, demonstrates this multiplicative form of value computation. Although this additional axiom is often violated in humans and monkeys (Allais, 1953; Starmer, 2000; Ferrari-Toniolo et al., 2022), a simple modification of the value computation mechanism employing subjectively weighted probabilities can account for these violations (Kahneman and Tversky, 1979; Camerer and Ho, 1994). Further work is necessary to establish whether amygdala neurons encode reward components compatible with this more reliable behavioral model, as recently reported in primate midbrain dopaminergic neurons encoding subjectively weighted reward probabilities (Ferrari-Toniolo et al., 2025).

How could the presently reported neurons coding subjective values contribute to broader amygdala functions? The behavioral relevance of value-coding neurons in amygdala may be reflected by effects of amygdala damage: in monkeys and humans, amygdala lesions alter reward-guided behaviors and economic decisions (Baxter and Murray, 2002; Brand et al., 2007; Murray, 2007; Levy et al., 2010; Machado et al., 2010; Talmi et al., 2010; van Honk et al., 2013; Costa et al., 2016; Rudebeck et al., 2017; Pujara et al., 2022). Consistently, human neuroimaging studies find subjective value signals in the amygdala (Levy et al., 2010; Grabenhorst et al., 2013; Zangemeister et al., 2016; Kim et al., 2024). In the present study, histological reconstructions showed that value-coding neurons were prevalent across the amygdala subdivisions we recorded from, including primarily the lateral and basolateral nuclei, but also (with fewer sampled neurons) in basomedial and central nuclei. As these nuclei differ in input and output connections, local circuit-designs and functions (Price et al., 1987; Pitkanen et al., 1997; Pitkanen and Amaral, 1998; Price, 2003; Sah et al., 2003; Maren and Quirk, 2004; Pape and Pare, 2010; Johansen et al., 2011; Duvarci and Pare, 2014; Gothard, 2020), value-coding neurons in each nucleus might contribute to different functions. The lateral nucleus might constitute the primary site for associating value information with visual cues (Johansen et al., 2011; Grabenhorst et al., 2019b) and distributing this information, via point-to-point connections (Pitkanen and Amaral, 1998), to the different downstream nuclei. The present results suggest that amygdala neurons encode values—including at this early processing stage—in a manner that reflects the subjective tradeoff between reward components. Previous studies showed value signals in lateral (but less so in basolateral) amygdala generalized across difference visual-cue formats (Grabenhorst and Baez-Mendoza, 2025) and across social self-other contexts (Grabenhorst et al., 2019b). Thus, subjective values compliant with EUT axiom encoded by lateral-nucleus neurons could be used flexibly for different downstream computations in amygdala. Different to the lateral nucleus, neurons in the basolateral amygdala have been more directly implicated in decision processes: basolateral neurons encode decision computations in an abstract, view-based representation (Grabenhorst et al., 2023), and integrate magnitude and probability information into risk (defined as the variance of expected rewards, distinct from value) (Grabenhorst and Baez-Mendoza, 2025). Thus, the presently reported value-coding neurons in basolateral amygdala might contribute to local decision computations and decision processes in other brain structures to which this nucleus projects, including the orbitofrontal cortex and striatum. For example, amygdala lesions have been shown to affect value-coding by neurons in the orbitofrontal cortex (Rudebeck et al., 2013; Rudebeck et al., 2017) and behavioral measures of value-updating (Murray and Rudebeck, 2013). Finally, value-coding neurons in central nucleus—the amygdala’s principal output structure to autonomic, endocrine, and attentional effector systems (Price, 2003; Johansen et al., 2011)—may play roles in translating value into physiological and emotional responses (Fustinana et al., 2021; Courtin et al., 2022).

The amygdala is not only a key reward structure involved in economic decisions but is also implicated in a variety of psychiatric and behavioral conditions in which reward valuation, reward-related decision-making, and mental health are affected (Oler et al., 2010; Price and Drevets, 2012; Bernardi and Salzman, 2019; Andrews et al., 2022; Klein-Flugge et al., 2022; Fox and Shackman, 2024). Thus, our present findings that amygdala neurons assign subjective values to visual cues, consistent with axioms of economic theory, and processes these values for decision-making may inform our understanding of the amygdala’s role in conditions that impair mental health and well-being. For example, reduced sensitivity of amygdala neurons to visually cued value information or deficient neuronal integration of value information may contribute to maladaptive preferences.

In conclusion, our results demonstrate compliance with the continuity axiom of EUT in both behavior and neuronal responses and behavioral-neuronal subjective value measures in the amygdala. These findings identify amygdala neurons as substrates for encoding behavior-matching subjective values according to the continuity axiom of EUT, and for translating these values into economic choices.

## ACKNOWLEDGEMENTS

We thank Aled David and Christina Thompson for animal care and technical support, Dr. Polly Taylor for expert anesthesia, Dr. Henri Bertrand for veterinary support, Dr. Raymundo Báez-Mendoza for expert contributions to animal training, surgery, and task programming. This work was funded by the Wellcome Trust and the Royal Society (Wellcome/Royal Society Sir Henry Dale Fellowship grants 206207/Z/17/Z and 206207/Z/17/A to F.G.; Wellcome Trust Principal Research Fellowship to WS, by Wellcome Grants WT 095495 and WT 204811 to W.S), the John Fell Oxford University Press Research Fund to F.G., and a Discovery Fellowship by the UK Biotechnology and Biological Sciences Research Council (BBSRC) to S.F.-T. This research was funded in whole, or in part, by the Wellcome Trust. For the purpose of Open Access, the author has applied a CC BY public copyright license to any Author Accepted Manuscript version arising from this submission.

## AUTHOR CONTRIBUTIONS

F.G. and W.S. designed research; F.G. performed research; F.G. and S.F.-T. analyzed data; F.G., W.S. and S.F.-T., wrote the paper.

